# DLK1 is a GATA1s-Driven Dependency and Therapeutic Target in Down Syndrome-Associated Myeloid Leukemia

**DOI:** 10.1101/2024.08.30.608947

**Authors:** Lonneke J. Verboon, Sonali P. Barwe, Meredith Tavenner, Joshua R. Faust, José Gonçalves-Dias, Konstantin Schuschel, Raj Bhayadia, Patrick H. van Berkel, Aimy Sebastian, Rhonda E Ries, Sophie Paczesny, Soheil Meshinchi, Johann Hitzler, Yana Pikman, E. Anders Kolb, Dirk Heckl, Jan-Henning Klusmann, Anilkumar Gopalakrishnapillai

## Abstract

Children with Down syndrome have a markedly increased risk of developing myeloid leukemia (ML-DS). Although having an excellent prognosis, 10–20% develop relapsed or refractory disease with poor survival, highlighting the need for new targeted approaches. The pathogenesis of ML-DS is tightly linked to fetal hematopoiesis and mutations in GATA1, generating the truncated GATA1short(s) isoform.

We identified Delta-like non-canonical Notch ligand 1 (DLK1) as a direct GATA1s target. DLK1, a paternally imprinted transmembrane protein, is highly expressed in fetal liver CD34⁺ cells but absent in adult hematopoiesis, making it an attractive immunotherapeutic target. Chromatin profiling revealed GATA1s occupancy at a distal enhancer within the DLK1–DIO3 locus, driving aberrant DLK1 upregulation in ML-DS. Functional studies demonstrated that DLK1 is a leukemia dependency, as its genetic ablation impaired proliferation and engraftment, induced apoptosis, and altered Notch and β-catenin signaling.

Therapeutically, a DLK1-directed antibody-drug conjugate (DLK1-ADC) induced selective cytotoxicity, abrogated colony formation, and significantly prolonged survival in refractory ML-DS PDX models, achieving durable remissions at higher doses. These findings establish DLK1 as a leukemia-specific vulnerability and provide preclinical proof-of-concept for DLK1-targeted therapies in ML-DS and other leukemias with fetal-like expression programs.

**Key Points:**

- DLK1 is a GATA1s-driven leukemia dependency in ML-DS, linking fetal hematopoietic programs to leukemic stemness and refractory disease.
- Therapeutic targeting of DLK1 with an antibody–drug conjugate selectively eradicates leukemia cells and prolongs survival in ML-DS PDX models

## Introduction

Children with Down syndrome (DS) have a striking predisposition to myeloid malignancies, with a ∼500-fold increased risk compared to the general population (1–5). The majority of cases arise in early childhood and are preceded by a preleukemic condition termed transient abnormal myelopoiesis (TAM), which is detected in up to 30% of neonates with DS (6). While TAM often resolves spontaneously, in 20–30% of children it progresses to myeloid leukemia of Down syndrome (ML-DS). Treatment of ML-DS with reduced-intensity chemotherapy has transformed outcomes, with current event-free survival rates approaching 90% (7–9). Nevertheless, DS patients particularly suffer from treatment related toxicity and mortality. Moreover, 10-20% of patients present with refractory disease or relapse early after therapy, and their survival drops below 25% (10). Effective therapeutic options for this group are lacking, highlighting the urgent need to identify new vulnerabilities that can be exploited in a targeted manner (11).

The biology of ML-DS is intimately linked to fetal hematopoiesis. Somatic mutations in *GATA1* – which generate the truncated transcription factor GATA1s – arise exclusively in fetal hematopoietic stem and progenitor cells (HSPCs) (12). These mutations alter transcriptional output leading to TAM (13). A defining feature of this fetal origin is the persistence of developmental gene expression programs that are normally extinguished after birth. Mounting evidence suggests that these fetal-restricted programs are central to leukemogenesis and may also represent selective therapeutic entry points, as they are absent from postnatal hematopoiesis (14, 15).

One such program involves Delta-like non-canonical Notch ligand 1 (*DLK1*). DLK1 is a paternally imprinted (16) transmembrane protein containing EGF-like repeats and is normally expressed at high levels in the fetal liver, kidney, and other developing tissues (17–19). In hematopoiesis, DLK1 marks immature HSPCs and contributes to the regulation of self-renewal and differentiation (20). Postnatally, DLK1 expression is strongly downregulated (21) and is essentially absent from normal adult CD34⁺ hematopoietic progenitors (22). Its re-expression has been documented in myelodysplastic syndromes (MDS), acute myeloid leukemia (AML) (22, 23), and several solid tumors (24), where it is associated with stemness, impaired differentiation, and inferior prognosis (25, 26). The restricted developmental expression pattern, together with negligible levels in normal adult hematopoiesis, makes DLK1 a highly attractive candidate for immunotherapeutic intervention. Preclinical strategies for DLK1 targeting have demonstrated anti-tumor activity, including antibody-drug conjugates (27, 28), chimeric antigen receptor (CAR) T cells in hepatocellular carcinoma models (29), and radioimmunotherapy with ^125^I-labeled anti-DLK1 antibodies in lung cancer (30).

Mechanistically, DLK1 acts as a non-canonical Notch ligand, capable of either suppressing or activating Notch receptor signaling in a context-dependent manner (25, 31). It can also signal independently through intracellular cleavage product that influences transcriptional programs (32). In myelodysplastic syndromes, elevated DLK1 expression has been linked to increased clonogenic growth and impaired differentiation of CD34⁺ progenitors, but its role and regulation in AML and ML-DS have remained largely undefined (22, 33).

In the present study, we identify *DLK1* as a direct transcriptional target of GATA1s in ML-DS. Chromatin profiling revealed GATA1s occupancy at a distal enhancer within the *DLK1–DIO3* locus, linking *GATA1s* mutations – a necessary initiating lesion in ML-DS – to aberrant *DLK1* upregulation. This establishes a mechanistic connection between fetal transcriptional programs and leukemic dependency. Building on this, we demonstrate that DLK1 can be therapeutically targeted using an antibody-drug conjugate (DLK1-ADC), which selectively eradicates DLK1-expressing cells and prolongs survival in refractory ML-DS xenograft models. These findings position DLK1 as a leukemia-specific vulnerability and provide preclinical proof-of-concept for DLK1-directed therapy in ML-DS and other leukemias with fetal-like expression programs.

## Results

### Aberrant DLK1 expression characterizes ML-DS

We first assessed *DLK1* expression in ML-DS by comparing transcriptomes of ML-DS cell/PDX lines (n=5) with normal bone marrow-derived CD34⁺ cells (n=4). *DLK1* emerged as one of the top upregulated surface proteins (**Supplementary Fig. S1A** and **Supplementary Table 1**). Analysis of pediatric AML datasets (AAML1531 and AAML08B1) confirmed significantly higher *DLK1* expression in ML-DS bone marrow samples (257.2 ± 73.8 TPM, n=77) compared with normal controls (2.2 ± 0.2 TPM, n=68) (**Fig. 1A**). In an independent cohort, *DLK1* expression was also enriched in ML-DS compared with normal adult peripheral blood stem/progenitor cells (HSPCs) and other cytogenetic subtypes of pediatric AML, including non-DS acute megakaryoblastic leukemia (AMKL; **Fig. 1B**) (34). Fetal-liver CD34^+^ HSPCs showed high DLK1 levels comparable to, and in some samples exceeding, those in TAM and ML-DS (34). No significant difference was observed between TAM and ML-DS samples in both datasets.

**Fig. 1.**
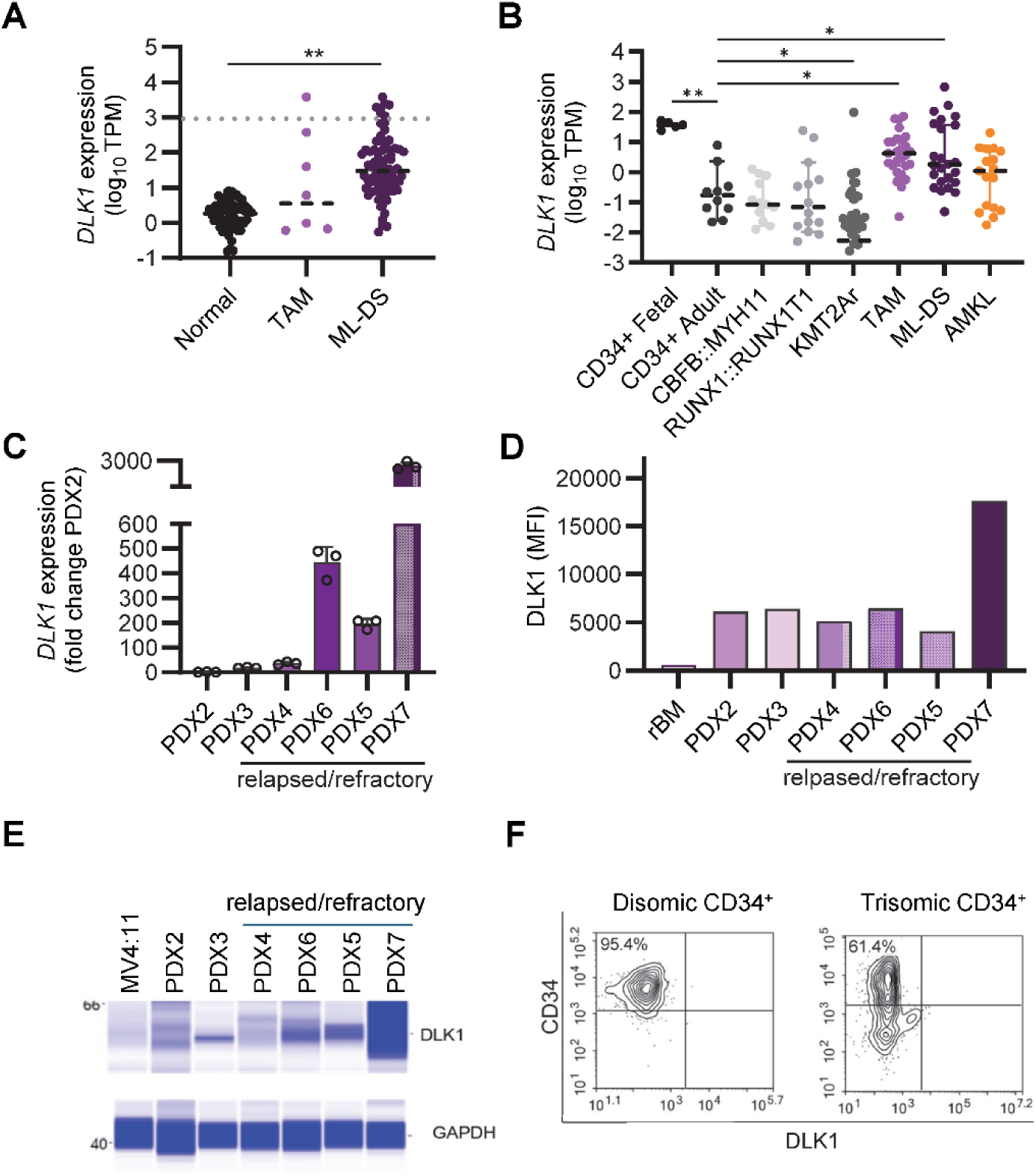
DLK1 expression in ML-DS. **A**) *DLK1* transcript levels in normal (n=68), TAM (n=8), and ML-DS (n=77) bone marrow specimens from AAML1531 and AAML08B1. **B**) *DLK1* expression (TPM) in fetal CD34^+^ (n=5) and adult CD34^+^ HSPCs (n=10), CBFB::MYH11 (n=13), RUNX1::RUNX1T1 (n=14), KMT2Ar (n=48), TAM (n=27), ML-DS (n=24), and AMKL (n=17). **C**) *DLK1* transcript level measured by qRT-PCR in ML-DS PDX lines. Fold change with respect to PDX2 is plotted. (n=3 mean ± SEM). **D**) DLK1 mean fluorescence intensities (MFI) in ML-DS PDX lines and in remission bone marrow (rBM) of one ML-DS patient. **E**) Immunoblots showing the expression of DLK1 in ML-DS PDX lines using Wes automated immunoblotting. GAPDH was used as a loading control. **F**) Representative flow cytometry plots of disomic (left) and trisomic (T21; right) CD34^+^ progenitors. Numbers indicate the percentage of events in the gated population (crosshair): 95.4% (disomic) and 61.4% (trisomic). Live singlets were pre-gated; fluorescence intensities are shown on a log scale. *p <0.05, **p <0.01

qRT-PCR confirmed higher *DLK1* transcript levels in PDX lines derived from refractory ML-DS compared with those from newly diagnosed cases (**Fig. 1C**). Flow cytometry and immunoblotting corroborated surface and protein expression in refractory PDXs, whereas DLK1 was undetectable in bone marrow from an ML-DS patient who went on to a achieve remission (rBM, **Fig. 1D-E**). Importantly, DLK1 was absent on CD34⁺ progenitors from individuals with or without DS (**Fig. 1F**), underscoring its disease-specific expression. Normal-tissue profiles from the Human Protein Atlas (35) show medium-high DLK1 protein in selected tissues (e.g. adrenal, placenta, ovary), with little or no detection in most other tissues including bone marrow (**Supplementary Fig. S1B)**, consistent with selective expression.

Together, these data establish DLK1 as a robustly expressed and selective marker of ML-DS, particularly enriched in refractory disease.

### GATA1s drives aberrant DLK1 expression through distal enhancer activation

To investigate why *DLK1* is aberrantly expressed in ML-DS, we characterized the epigenetic landscape of the locus. Chromatin profiling demonstrated that the *DLK1* promoter carries a bivalent signature (H3K4me3 and H3K27me3) in CD34⁺ cells and is enriched for H3K4me3 in megakaryocytes (**Fig. 2A**), consistent with active transcription (**Supplementary Fig. S2A**) and the previously described activity of the *DLK1-DIO3* locus in the megakaryocytic lineage (22). In contrast, the promoter displays repressive H3K27me3 in monocytes. In the ML-DS cell line CMK, Cleavage Under Targets & Release Using Nuclease (CUT&RUN) using exogenous HA-tagged GATA1s identified GATA1s occupancy at a distal enhancer upstream of DLK1, coinciding with H3K4me3 and H3K27ac, marks of active transcriptional programs. This enhancer lies within the imprinted *DLK1-DIO3* locus, which also harbors the non-coding RNA *MEG3*. To confirm that GATA1s also binds to this enhancer *in vivo*, we performed CUT&TAG for H3K27ac, CUT&RUN for GATA1s, and ATAC-seq on one TAM and one ML-DS patient sample, which showed a similar GATA1s occupancy pattern (**Fig. 2A**). Targeted CRISPR mutagenesis of two enhancer peaks (A and B) centered on the GATA1s site significantly reduced expression of both *DLK1* and *MEG3* in CMK cells (**Fig. 2B**), establishing enhancer–promoter cooperation within this domain.

**Fig. 2.**
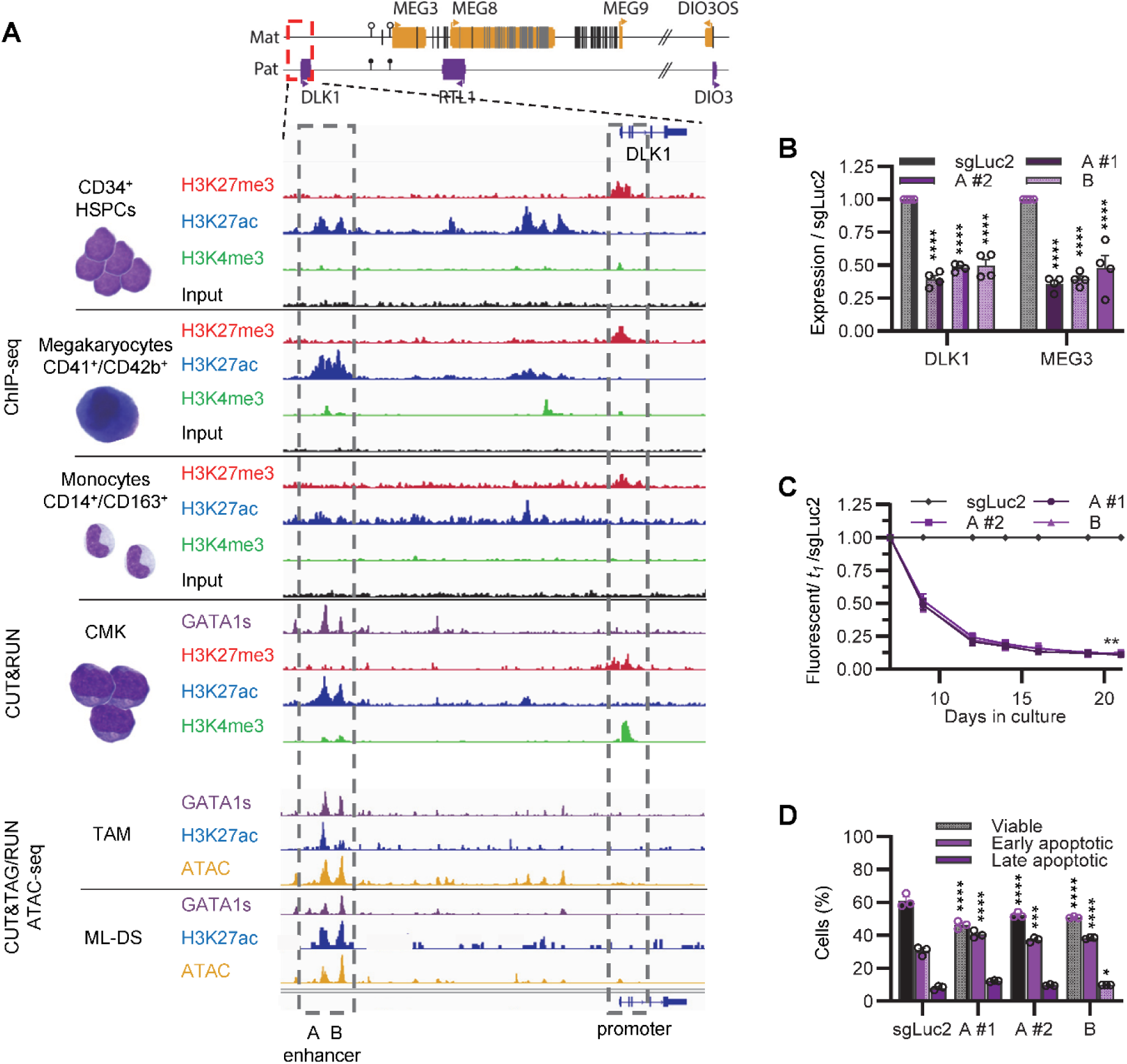
Distal enhancer regulation of DLK1 at the DLK1-DIO3 locus. **A**) Schematic of the DLK1-DIO3 locus (top) and tracks for H3K237me3 (red), H3K27ac (blue), H3K4me3 (green), input (black) in CD34+ PB-HSPCs, megakaryocytes (CD41+/CD42b+), and monocytes (CD14+/CD163+) (ChIP-seq), plus GATA1s-HA (dark purple) in CMK (CUT&RUN). GATA1s (dark purple, CUT&RUN), H3K27ac (blue, CUT&TAG), and ATAC-seq (Orange) in TAM and ML-DS. Grey box marks distal enhancer with peak A (marked A) and peak B (marked B) upstream of DLK1. **B**) qRT-PCR of *DLK1* and *MEG3* after CRISPR targeting peak A (two sgRNAs) or B (one sgRNA), shown relative to sgLuc2 (Normalized to B2M, n = 4 biological replicates, mean ± SEM). **C**) Proliferation of sgRNA+ (fluorescent) cells after enhancer targeting (n = 5 biological replicates, mean ± SEM; data normalized to sgLuc2 at t1 = 100%). Where only one set of asterisk is shown, all conditions share the same p value. **D**) Annexin V assay (early/late apoptosis) after enhancer targeting versus sgLuc2 (n = 3 biological replicates, mean ± SEM). **p <0.01

These findings demonstrate that GATA1s directly drives aberrant *DLK1* expression through distal enhancer activation at the *DLK1-DIO3* locus, mechanistically linking the initiating *GATA1s* mutation of ML-DS to reactivation of a fetal stemness regulator.

### DLK1 maintains leukemic proliferation and engraftment

To assess whether DLK1 contributes functionally to leukemic growth, we performed genetic perturbation in the ML-DS cell line CMK. Perturbing *DLK1* and *MEG3* expression using CRISPR mutagenesis of two enhancer peaks led to a significant growth disadvantage and induction of apoptosis (**Fig. 2C-D**). Similarly, CRISPR/Cas9-mediated knockout of *DLK1* using two independent sgRNAs efficiently reduced DLK1 surface expression (**Fig. 3A**) and resulted in progressive loss of DLK1-deficient cells in competitive proliferation assays (**Fig. 3B**). This depletion was accompanied by increased apoptosis (**Fig. 3C**). Still, the effects of *DLK1* knockout were not as strong as perturbing the enhancer, which might be explained by additional effects mediated by the depletion of *MEG3*. Importantly, independent single-cell *DLK1* knockout clones (**Supplementary Fig. S2B**) further confirmed a strong proliferative defect, with a >60% reduction in EdU incorporation compared to parental CMK cells (**Fig. 3D**). Partial suppression by shRNA knockdown also impaired cell growth, demonstrating a graded dependency on *DLK1* expression.

**Fig. 3.**
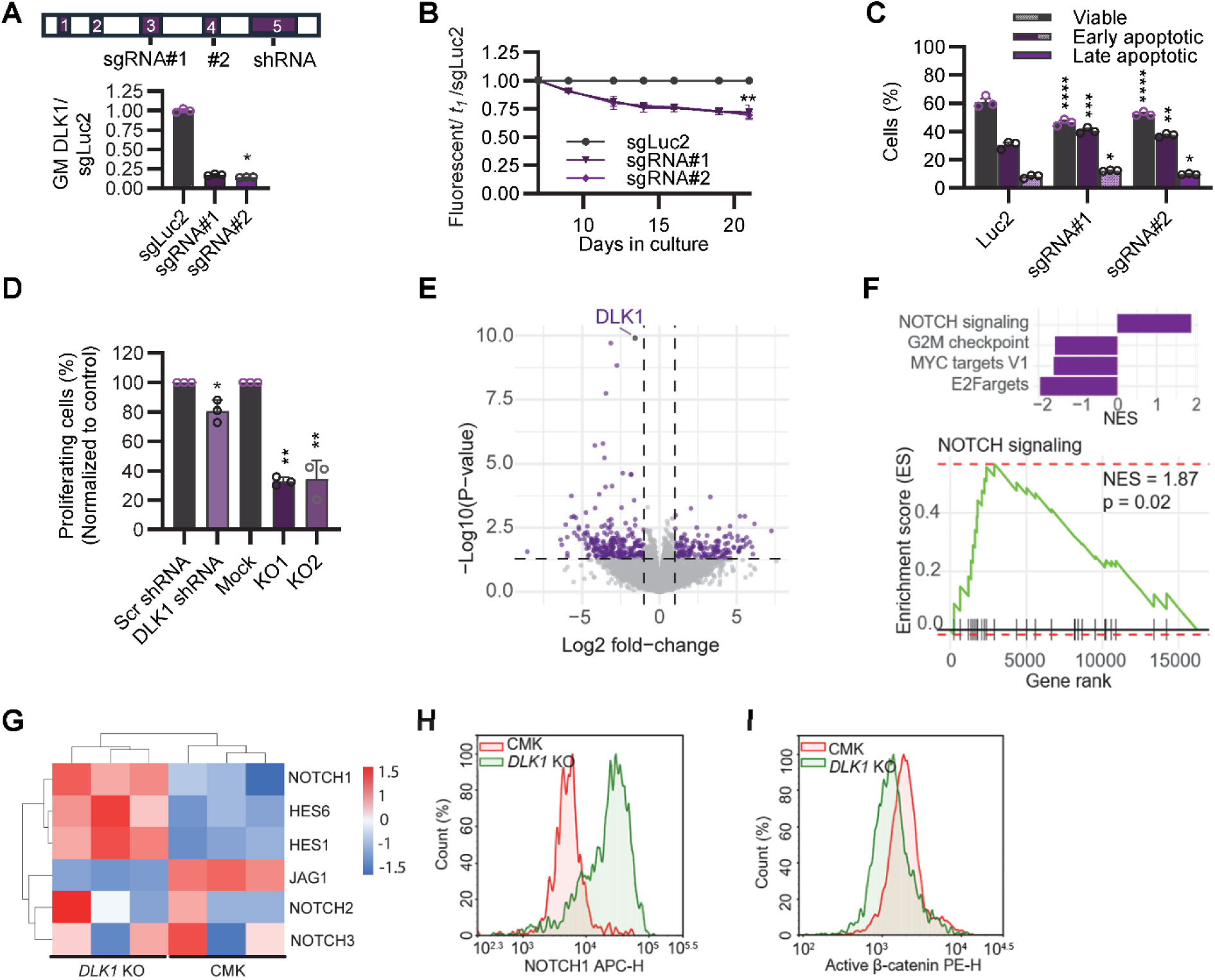
DLK1 perturbation reduces proliferation and shifts pathway activity. **A**) Graphical overview of the DLK1 gene depicting sgRNA and shRNA locations (top) and DLK1 surface levels (APC; geometric mean fluorescence; bottom) normalized to sgLuc2 (n = 3 biological replicates, mean ± SEM). **B**) Proliferation assay of sgRNA-positive (fluorescent) cells after targeting of DLK1 (n = 3 biological replicates, mean ± SEM; data normalized to sgLuc2 at t1 = 100%). Where only one set of asterisks is shown, all conditions share the same p value). **C**) Annexin V assay (early/late apoptosis) after targeting DLK1 versus sgLuc2 (n = 3 biological replicates, mean ± SEM). **D**) EdU assay of proliferating cells (%), normalized to control (n = 3 biological replicates, mean ± SEM). **E**) Volcano plot of bulk RNA-seq (sgRNA#2 vs sgLuc2). Dashed lines mark log_2_FC = 1 and p = 0.05; DLK1 is highlighted. **F**) Hallmark gene-set enrichment (MSigDB) for the same contrast. Bar plot shows significantly enriched pathways (FDR q = 0.05); bottom, GSEA enrichment plot for Notch signaling (NES = 1.87, p = 0.02). **G**) Heatmap of selected Notch pathway genes comparing *DLK1* KO and CMK. **H**) Notch1 cell surface protein levels (%) by flow cytometry (APC-H) in *DLK1* KO versus CMK. **I**) Active β-catenin by flow cytometry (PE-H) *DLK1* KO versus CMK. *p <0.05, **p <0.01

To explore the underlying mechanism, we performed bulk RNA-seq of sgRNA-transduced CMK cells. Loss of *DLK1* resulted in broad transcriptional changes, with 406 genes differentially expressed. Gene set enrichment analysis highlighted significant activation of the Notch pathway (**Fig. 3E-F**). Specifically, in single-cell *DLK1*-knockout clones *NOTCH1* and its downstream targets *HES1* and *HES6* were upregulated, while canonical Notch ligand *JAG1*, as well as *NOTCH2* and *NOTCH3*, were downregulated (**Fig. 3G**). Consistent with these findings, flow cytometry confirmed increased NOTCH1 protein expression and reduced levels of active β-catenin (**Fig. 3H-I**), suggesting that DLK1 sustains leukemic proliferation by suppressing NOTCH1 activity and maintaining β-catenin signaling.

We next evaluated the role of *DLK1* in leukemic propagation *in vivo*. Immunodeficient recipient mice transplanted with single-cell-derived *DLK1*-knockout CMK cells displayed dramatically reduced engraftment: human blasts were virtually absent in bone marrow and spleen compared to mice receiving wild-type CMK cells, which developed extensive infiltration and tissue destruction (**Fig. 4A-B**). Median survival was significantly prolonged in recipients of both sexes, with *DLK1*-knockout cohorts living more than 60 days longer than controls (**Fig. 4C**). Competitive homing assays further revealed impaired early localization of *DLK1*-deficient cells to bone marrow niches (**Fig. 4D**).

**Fig. 4.**
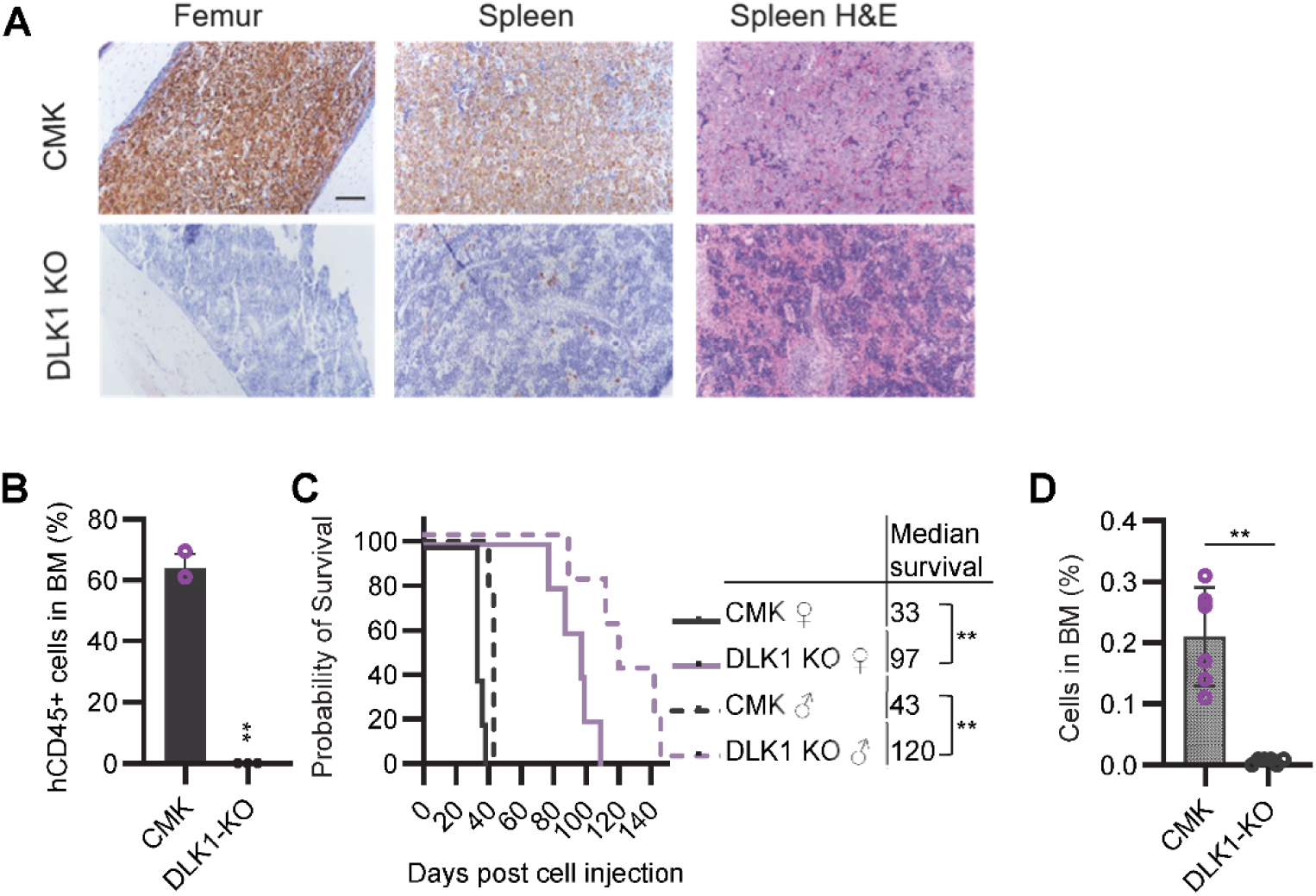
CMK cells with DLK1 knockout show substantial reduction in engraftment. **A**) Immunohistochemistry showing the presence of human cells (detected by staining with anti-human mitochondrial antibody). Hematoxylin and eosin (H & E) staining of spleen sections. Representative images from n=3 each. Bar = 500 um. **B**) CMK or *DLK1* KO cell percentage detected by flow cytometry in bone marrow cells harvested by flushing femurs following euthanasia (n=3 each). **C**) Kaplan-Meier survival estimates of male or female mice injected with CMK WT or *DLK1* knockout (KO) cells. n=5 per group. **D**) CMK *DLK1* KO cells found in the bone marrow in a competitive homing assay. n=6 per group *p<0.05, **p <0.01, ***P<0.001.

Together, these results demonstrate that *DLK1* is not merely a marker but a functional dependency of ML-DS, required for proliferation, survival, homing, and long-term engraftment. By sustaining leukemic stemness programs through modulation of Notch and β-catenin pathways, DLK1 acts as a key driver of disease maintenance *in vivo*.

### Therapeutic targeting of DLK1 with an antibody-drug conjugate

Given the essential role of *DLK1* in sustaining leukemic growth, we next evaluated its therapeutic potential using a DLK1-directed antibody-drug conjugate (DLK1-ADC; ADCT-701), consisting of an anti-DLK1 antibody conjugated to the pyrrolobenzodiazepine dimer payload PL1601 (28, 36, 37). *In vitro*, DLK1-ADC induced potent and selective cytotoxicity in CMK cells, with an IC_50_ of 0.027 nM, compared to 0.172 nM in *DLK1* knockdown cells and complete resistance in *DLK1* knockout cells (**Fig. 5A**). These results demonstrate strict dependence of ADC activity on DLK1 surface expression. Isotype control ADCs had no effect, confirming target specificity.

**Fig. 5.**
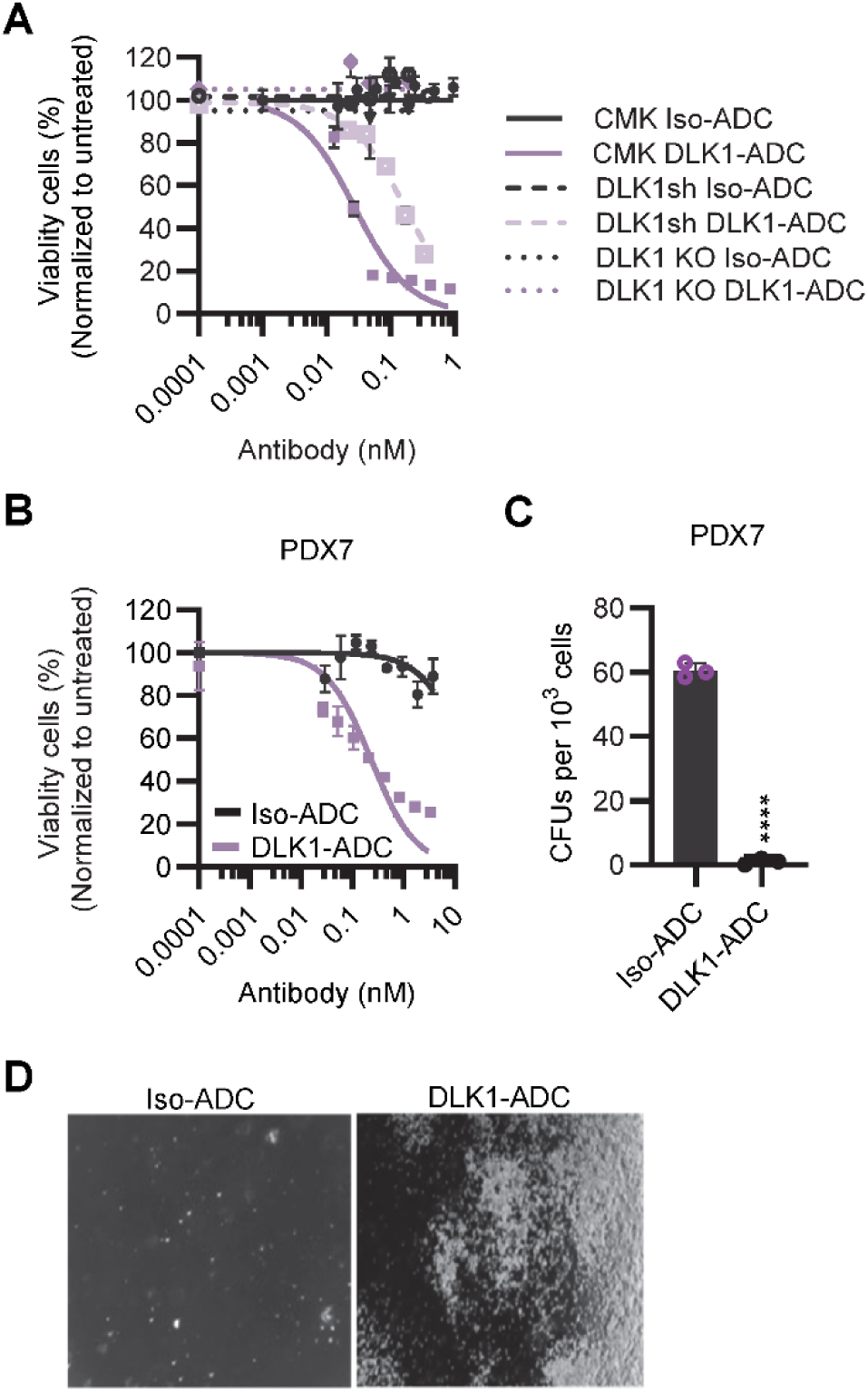
Targeting DLK1 using antibody drug conjugate ADCT-701. **A**) *In vitro* viability assay in CMK wild type (WT), CMK *DLK1* shRNA knockdown (DLK1sh) and CMK *DLK1* knockout (*DLK1* KO) cells. n = 3 biological replicates. **B**) Viability of PDX7 cells treated with varying concentrations of isotype ADC (Iso-ADC) or DLK1-ADC. **C**) Effect of DLK1-ADC (2.67 nM) on PDX7 colony formation in Methocult. **D)** Representative images are shown. Scale bar =100 um. n = 3 biological replicates. ****p<0.0001.

We extended these findings to primary ML-DS PDX samples. In refractory PDX7, which expressed high levels of DLK1, DLK1-ADC showed nanomolar potency (IC_50_ 0.234 nM) and completely abrogated colony formation in methylcellulose assays (**Fig. 5B-D**).

*In vivo*, DLK1-ADC significantly reduced leukemia burden and prolonged survival in PDX-engrafted recipient mice. A single dose of 0.25-1 mg/kg led to dose-dependent improvements in median survival, extending life expectancy by up to 24 days compared to isotype controls (**Fig. 6A, B**). Across three independent refractory ML-DS PDX models, repeated dosing at 1 mg/kg consistently prolonged survival, with the strongest effect observed in high-DLK1 expressers (**Fig. 7A, B**). Remarkably, a single 3 mg/kg dose in PDX5 induced durable remissions in two of three mice, with no evidence of disease recurrence (**Fig. 7C, D**).

**Fig. 6.**
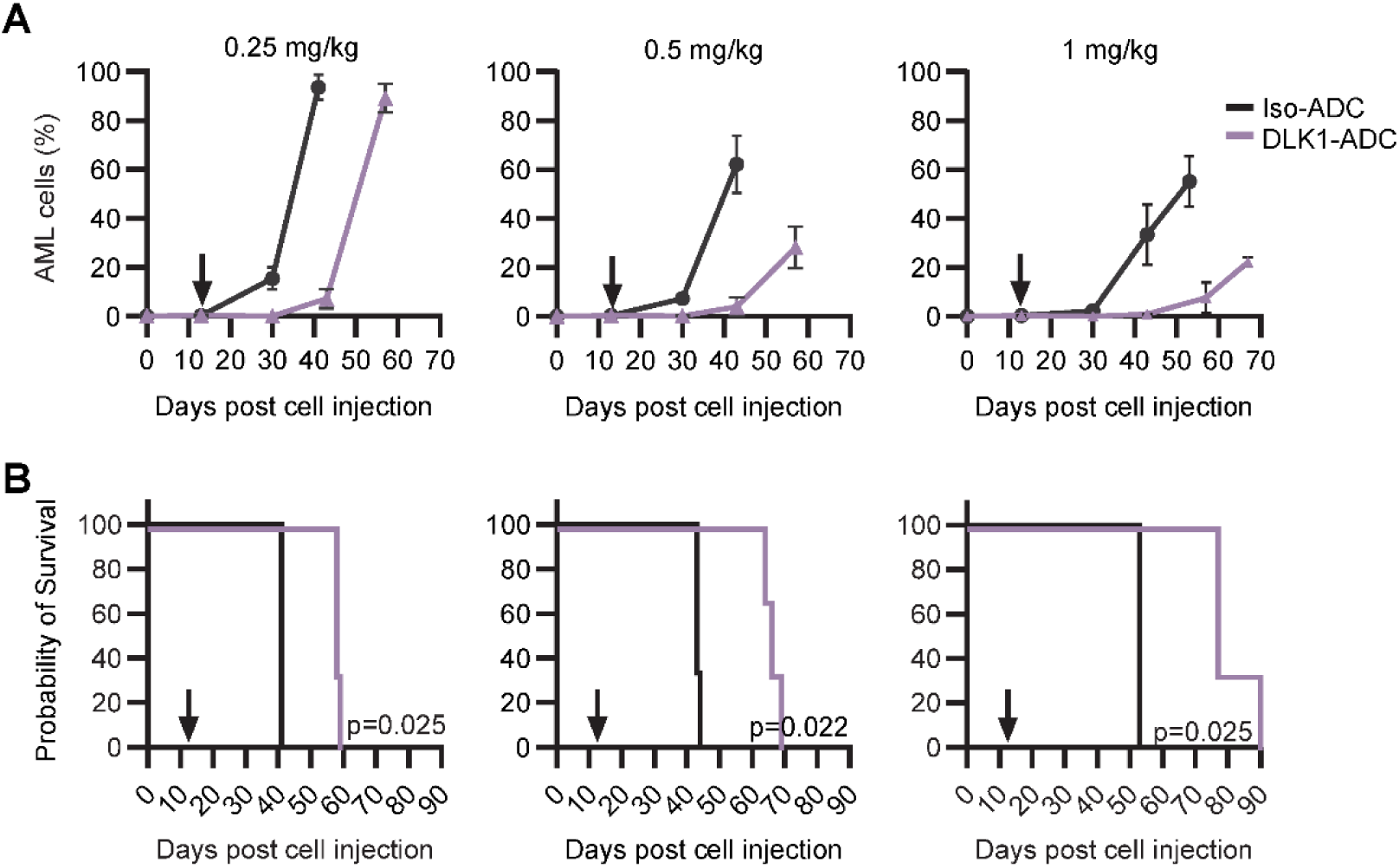
*In vivo* evaluation of antibody drug conjugate ADCT-701 in PDX7. **A**) Growth curve showing the rise in the percentage of AML cells in peripheral blood in mice transplanted with PDX7 and treated with DLK1-targeting ADCT-701 (DLK1-ADC) or with isotype control antibody drug conjugate B12-PL1601 (Iso-ADC) at indicated doses. **B**) Kaplan-Meier survival estimates of mice treated with ADCT-701 (DLK1-ADC) or B12-PL1601 (Iso-ADC). Arrows indicate time when ADC was administered intravenously. n=3 per group, *p <0.01.

**Fig. 7.**
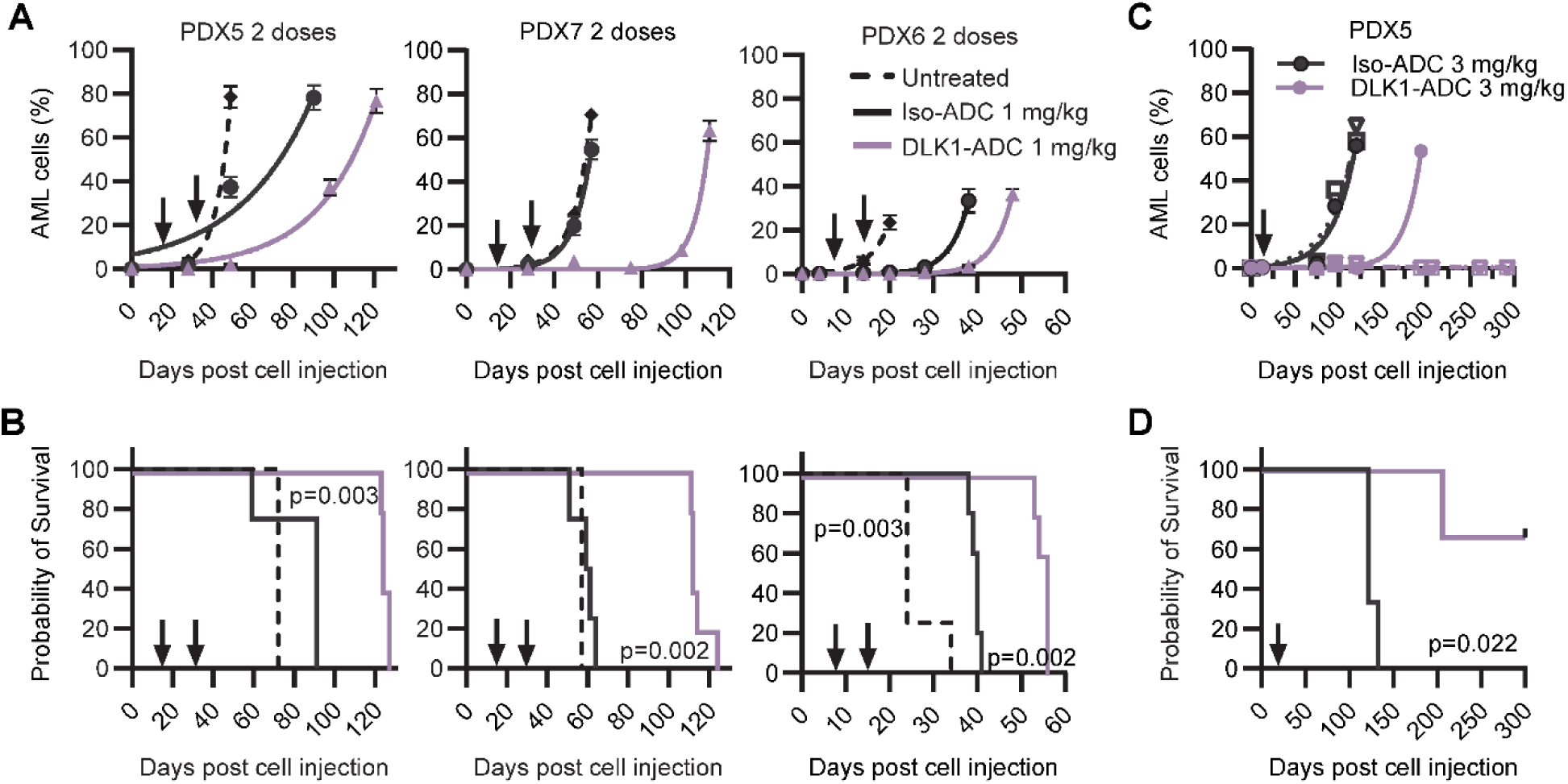
*In vivo* evaluation of antibody drug conjugate ADCT-701 in ML-DS PDX models. **A**) Growth curve showing the rise in the percentage of AML cells in peripheral blood in mice transplanted with PDX5, PDX7, and PDX6 respectively. **B**) Kaplan-Meier survival estimates of mice treated with DLK1-targeting ADCT-701 (DLK1-ADC) or isotype control antibody drug conjugate B12-PL1601 (Iso-ADC). Arrows indicate time when ADC was administered intravenously. n=4-5 per group. **C**) Growth curve showing the rise in the percentage of AML cells in peripheral blood in mice transplanted with PDX5. Note, that each line represents individual mouse. **D**) Kaplan-Meier survival estimate of mice treated with ADCT-701 (DLK1-ADC) or B12-PL1601 (Iso-ADC). Arrows indicate time when ADC was administered intravenously. *p <0.01, n=3 per group.

These data demonstrate that DLK1-ADC selectively eradicates DLK1-positive leukemic cells and achieves meaningful survival benefit *in vivo*. The correlation between surface expression and therapeutic efficacy further highlights DLK1 as a tractable immunotherapeutic target in ML-DS.

## Discussion

Despite excellent survival for most children with ML-DS treated with reduced-intensity chemotherapy, relapsed or refractory disease remains a major challenge, with survival rates below 25% (10). Our study identifies DLK1 as a mechanistic driver and a tractable vulnerability in ML-DS. By tying aberrant DLK1 expression to GATA1s-driven enhancer activation within the *DLK1-DIO3* locus, our study provides a rationale and preclinical proof-of-concept for therapeutically targeting this fetal stemness regulator that is normally silenced in adult hematopoiesis.

Expression analyses revealed that *DLK1* is strongly upregulated in ML-DS compared to normal hematopoietic progenitors, and is particularly enriched in refractory disease, while absent from CD34⁺ progenitors of healthy individuals. This finding is consistent with earlier observations that *DLK1* expression is largely restricted to fetal hematopoiesis and development, but re-emerges in a range of malignancies including MDS, AML (22, 23, 38), neuroblastoma (27), and hepatocellular carcinoma (25, 39). In mouse models, *DLK1* is expressed in the aorta-gonad-mesonephros region where it regulates emerging HSPCs by restricting their expansion through membrane-bound DLK1 rather than soluble isoforms (40). Moreover, in mice the *Dlk1-Gtl2* imprinted locus, which also harbors *MEG3*, is required for long-term HSC maintenance by modulating mitochondrial metabolism and limiting oxidative stress (41). These developmental roles illustrate how DLK1 sustains stemness in fetal contexts, explaining why its dysregulation in ML-DS enforces a fetal-like leukemic state and is directly coupled to disease pathogenesis.

Mechanistically, we show that *DLK1* is a transcriptional target of GATA1s, product of the mandatory initiating mutation in ML-DS. Chromatin profiling revealed GATA1s occupancy at a distal enhancer within the *DLK1-DIO3* locus, establishing a direct connection between GATA1s-driven transcriptional control and aberrant *DLK1* upregulation. This enhancer-promoter cooperation also impacted expression of *MEG3*, a non-coding RNA implicated in hematopoietic regulation, (42) consistent with reports that the *DLK1-DIO3* imprinted domain acts as a hub of developmental gene regulation and stem cell maintenance (41). In ML-DS, *GATA1s* mutations hijack this fetal circuitry to enforce *DLK1* expression and leukemic dependency These findings underscore how developmental programs normally extinguished after birth are exploited in ML-DS to sustain leukemic growth.

Functional studies confirmed that *DLK1* is not merely a marker but a critical dependency. Genetic ablation reduced proliferation, induced apoptosis, and profoundly impaired *in vivo* engraftment. Transcriptomic profiling of *DLK1*-deficient cells revealed activation of NOTCH1 and suppression of β-catenin signaling, indicating that DLK1 sustains leukemic stemness by balancing developmental pathways. DLK1’s dual role as a non-canonical Notch ligand (31, 43–46) and regulator of Wnt/β-catenin signaling has been described in other contexts, including hepatocellular carcinoma and glioblastoma, where it supports tumor initiation and progression (47). In normal hematopoiesis, *DLK1* expression in fetal liver HSPCs has also been linked to enhanced self-renewal capacity. Together, these data highlight DLK1 as a nexus of fetal transcriptional programs and leukemic stemness.

*DLK1* thus represents a leukemia-specific dependency directly linked to the ontogeny of ML-DS and to the obligate *GATA1s* mutation. The therapeutic implications of these findings are considerable. Unlike broadly expressed AML antigens such as CD123, CD33, or CLL-1, *DLK1* is absent from adult hematopoiesis, making it a promising target with reduced risk of myelosuppression. Its surface localization and high density in ML-DS further enhance its suitability for antibody-based strategies. Our preclinical testing of a DLK1-directed antibody-drug conjugate (ADCT-701) demonstrated potent, expression-dependent activity: nanomolar cytotoxicity in cell lines and PDXs, elimination of colony-forming capacity, and significant survival benefits *in vivo*. Notably, durable remissions were achieved at higher doses in refractory PDX models, providing compelling evidence that DLK1 can be therapeutically exploited. These data align with growing evidence that DLK1 is a tractable immunotherapeutic target across cancers with activated or reactivated oncofetal expression programs. In hepatocellular carcinoma, DLK1-directed CAR-T cells achieved strong anti-tumor activity in preclinical models (29). In lung cancer, ^125^I-labeled anti-DLK1 antibodies demonstrated efficacy in radioimmunotherapy studies (30). A DLK1-ADC has also been tested in neuroblastoma and adrenocortical carcinoma, showing selective killing of DLK1-expressing tumors (36, 37). Together, these studies highlight the translational feasibility of DLK1-directed therapies.

Our findings further highlight the broader concept that developmental dependencies can be leveraged as therapeutic vulnerabilities in pediatric leukemia. In recent years, fetal-restricted programs have been implicated in infant AML subtypes such as *NUP98*- or *KMT2A*-rearranged leukemia, where persistence of embryonic transcriptional networks enforces stemness and therapy resistance (48, 49). Targeting such fetal programs – whether through epigenetic modulators or surface antigens like DLK1 – offers a rational strategy to improve outcomes in diseases where conventional cytotoxic intensification is not feasible or inefficient. The demonstration that GATA1s, the founding lesion of ML-DS, directly drives *DLK1* expression places *DLK1* within this conceptual framework and strengthens its appeal as a rational therapeutic target.

Some limitations must be considered. Our PDX analyses, while robust, were limited in number; larger cohorts will be required to confirm its clinical utility. Potential toxicities remain to be evaluated, especially in infants or regenerative contexts where residual *DLK1* expression might persist. Mechanistically, the relative contributions of Notch-dependent versus Notch-independent pathways downstream of DLK1, as well as potential functions of soluble DLK1 isoforms, require further investigation.

Nevertheless, our results establish DLK1 as a leukemia-specific dependency in ML-DS, directly tied to its fetal origin and GATA1s-driven enhancer activation. By integrating developmental biology with therapeutic targeting, we provide both mechanistic insight and translational promise. DLK1’s restricted expression pattern, strong disease association, and amenability to multiple therapeutic modalities underscore its appeal as a target. These findings support clinical development of DLK1-directed therapies to improve outcomes for children with refractory ML-DS and potentially for other pediatric leukemias with fetal-like expression programs.

## Materials and Methods

### Cell culture

HEK293T and CMK (ACC 392) cells were obtained from DSMZ (DSMZ -German Collection of Microorganisms and Cell Cultures GmbH) and cultured according to the supplier’s recommendations and as described previously (14, 50). For culturing PDX cells, StemSpan SFEM media (StemCell Technologies, Ontario, Canada) supplemented with TPO 25ng/mL, IL3 10 ng/mL and GM-CSF 10 ng/mL (from R&D Systems, Minneapolis, MN) was used. Human CD34^+^ hematopoietic stem and progenitor cells (HSPCs) were isolated from mobilized peripheral blood from anonymous healthy donors and enriched using CD34^+^ immunomagnetic microbeads (Miltenyi Biotech, Bergisch Gladbach, Germany). Human HSPCs were expanded in StemSpan SFEM with 1% penicillin/streptomycin, 50 ng/mL SCF, 50 ng/mL FLT3-ligand, 20 ng/mL TPO, 10 ng/mL IL-3, and 10 ng/mL IL-6. For megakaryocyte differentiation, cells were cultured in StemSpan SFEM with 1% penicillin/streptomycin, 0.25% CD lipid concentrate (Gibco, Thermo Fisher Scientific, Waltham, MA), 20 ng/mL TPO, 13.5 ng/mL IL-9, 7.5 ng/mL IL-6, and 1 ng/mL SCF. For monocyte differentiation, cells were cultured in RPMI 1640 Glutamax, 10% FCS, 1% penicillin/streptomycin, 50 ng/mL FLT3 ligand, 50 ng/mL M-CSF, 10 ng/mL GM-CSF, 10 ng/mL IL-3, and 10 ng/mL IL-6. On day 7, FLT3 ligand and IL-3 were removed from the media. All human cytokines were purchased from Peprotech (Rocky Hill, NJ).

### RNA-Sequencing

Four ML-DS PDX lines, CMK, and primary normal human bone marrow-derived CD34^+^ cells from ATCC (American Type Culture Collection, PCS-800-012), and *DLK1* KO CMK cells were used for total RNA isolation using RNeasy Plus Micro kit (Qiagen, Germantown, MD). RNA sequencing library preparations, sequencing reactions, and preliminary bioinformatics analysis was conducted at GENEWIZ, LLC (South Plainfield, NJ). RNA-Sequencing was performed on 18 samples, 5 samples of ML-DS and 4 samples of normal cells in duplicates to generate 150 bp long paired-end reads. RNA-Seq analysis was performed as described previously (51), by aligning reads to hg38 using STAR aligner (52) and summarizing read counts per gene locus using featureCounts (53). Differentially expressed genes were identified using edgeR (54). UniProt list of membrane-associated proteins based on subcellular locations was used for filtering cell surface proteins. Similar analysis pipeline was used for single cell *DLK1* KO clones.

RNA-Seq data from bone marrow specimens of normal individuals, and patients with TAM or ML-DS were obtained from the TARGET (Therapeutically Applicable Research to Generate Effective Treatments): Acute Myeloid Leukemia (AML) database, study ID phs000465.v23.p8.

For *DLK1* KO samples, sorting was performed on day 3, and strand-specific eukaryotic mRNA (polyA enriched) total cellular RNA sequencing was performed by Novogene GmbH on an Illumina NovaSeq X plus using 150 bp paired-end chemistry. The received raw fastq files were processed using the nf-core/rna pipeline (v. 3.12.3) (55) and the quality was checked using fastQC (56). The adapters were trimmed with Trimgalore (57) and aligned against human genome hg38 using STAR aligner (52), and post gene quantification was determined with salmon (58). Differentially expressed genes were identified using edgeR (54). Gene set enrichment analysis for the gene set Hallmark was performed in R using the msigdr (v. 24.1.0) package (59).

### Quantitative real time polymerase chase reaction (qRT-PCR)

Total RNA was isolated using RNeasy Plus Micro kit following manufacturer’s protocol. cDNA was amplified from 1 mg total RNA using iScript^TM^ cDNA Synthesis kit (BioRad, Hercules, CA). qRT-PCR reaction was performed using PowerUp^TM^ SYBR^TM^ Green Master Mix or TaqMan™ Fast Advanced Master Mix (Applied Biosystems, Waltham, MA) for qPCR in a QuantStudio 7 Real-Time PCR (Polymerase Chain Reaction) System (Thermo Fisher Scientific). The relative quantification method using delta delta Ct calculation was used to determine the fold change in transcript levels. When a gene was not expressed the Ct value was set on 40. Primer sequences and Taqman assay number are listed in **Supplementary Table 2**.

### Flow cytometry

Cells were resuspended in phosphate-buffered saline containing 1% FBS and stained in the dark for 15 min with APC-conjugated anti DLK1 antibody (R & D Systems, #FAB1144A) or APC-conjugated Notch1 antibody (BioLegend, San Diego, CA, #352107). For intracellular flow cytometry, cells were fixed with fixation buffer, permeabilized with 90% methanol and stained with PE-conjugated non-phospho (active) β-catenin (Cell Signaling Technology, clone D13A1) in cell staining buffer for 1 hour. Samples were run on a Novocyte Quanteon flow cytometer (Agilent Technologies, Santa Clara, CA). Mouse blood was stained using CD45 mouse and human antibodies (BioLegend) and run on flow cytometer following RBC lysis. The following antibodies were used for sorting megakaryocytes: CD41:PE-Cy7 (Beckman Coulter, Brea, CA), CD61:PE-CY7 (Beckman Coulter), and CD42b:APC (BD, Franklin Lakes, NJ) and for monocytes: CD14:APC (Beckman Coulter), and CD163:PE (BD Biosciences, San Jose, CA). The following antibodies were used for sorting patient leukemic blasts: CD33:FITC, CD45:APC, CD3:APC-Cy7, CD7:APC-H7, CD19:APC-Cy7 (all BD Pharmingen, BD Biosciences) Apoptosis was measured using the Annexin V Apoptosis Detection Kit II with APC-Annexin V (BD Biosciences).

### Fluorescence-based proliferation assays

Individual proliferation assays were conducted using stable Cas9-expressing cell lines for CRISPR/Cas9 experiments as described previously (14, 60). Cells were transduced with individual sgRNA constructs at an efficiency of ∼40% to attain a mixed population, allowing for direct competition between transduced and untransduced cells. These cultures were maintained for up to 21 days, during which fluorescence was tracked every 2–3 days via flow cytometry. Depletion curves were generated by normalizing the percentage of fluorescent (i.e., transduced) cells at each time point to both the initial fluorescence (day 7) and the non-targeting control (sgLuc2).

### Western blot

Automated immunoblot analysis was performed using the Wes system (ProteinSimple, San Jose, CA) following manufacturer’s instructions using a 12-230 kDa Separation Module (SM-W001) and the anti-rabbit Detection Module (DM-001) as described previously (51). Cells were lysed in Minute Total Protein Extraction kit (Invent Biotechnologies, Plymouth, MN), sonicated and clarified via centrifugation 16000 x g for 15 min. The supernatant was collected and loaded into the capillary based on the number of cells lysed. Antibodies were purchased from Cell Signaling Technology (Danvers, MA). Normalization to total protein was performed using the Total Protein Detection Module in Wes (DM-TP0010).

### Lentiviral vectors and transduction

Individual single guide RNAs (sgRNAs) for *DLK1* were designed using CCTop (https://cctop.cos.uni-heidelberg.de/) and cloned via Esp3I into the SGL40C.EFS.dTomato (Addgene 89395) as described previously (60). Non-targeting control sgRNAs were designed to target the firefly luciferase gene.

Lentiviral particles were produced by co-transfection of the expression vector and packaging plasmid pMD2.G and psPAX2 (Addgene 12259 and 12260, respectively) into HEK293T cells using polyethylenimine (PEI; Polysciences, Warrington, PA). The viral particles were concentrated via high-speed centrifugation. Transduction was performed in normal cell culture medium. sgRNA sequences are listed in **Supplementary Table 3.**

### CRISPR genome editing (RNP nucleofection) and shRNA knockdown

CRISPR guide sequence for DLK1 was chosen from one of the pre-optimized sequences provided by IDT (Coralville, IA). Alt-R CRISPR-Cas9 sgRNA with single RNA molecule comprising crRNA and tracrRNA was complexed with Alt-R Cas9-GFP nuclease (IDT) to form RNP complex, which was introduced into CMK cells using a 4D nucleofector system and P3 primary cell 4D-Nucleotransfector X Kit L (Lonza, Basel, Switzerland). Post transfection, cells were cultured in 12-well plates. Individual wells were checked for DLK1 expression by flow cytometry. Clones with DLK1 expression overlapping with isotype control antibody were expanded and used for TIDE analysis (see below).

For DLK1 knockdown, pLV[shRNA]-EGFP:T2A:Puro-U6>DLK1 (VectorBuilder, Neu-Isenburg, Germany) shRNA lentiviral vector was used to generate lentiviral particles. CMK cells were spinoculated by centrifugation. Cells were cultured in puromycin to enrich the transduced cells. sgRNA and shRNA sequences and are listed in **Supplementary Table 3.**

### CRISPR editing efficiency (TIDE)

Genomic DNA was isolated with using a MicroDNA kit (Qiagen, Germantown, MD) or a Quick-DNA Miniprep Kit (Zymo Research Europe, Freiburg, Germany). The target locus was PCR-amplified with Q5® High-Fidelity DNA Polymerase (New England Biolabs, Frankfurt am Main, Germany) using primers flanking the expected Cas9 cut site region surrounding the CRISPR cut site in the genomic DNA. PCR products were Sanger sequenced and analyzed using a web-based tool https://ice.synthego.com/ (https://ice.synthego.com or https://apps.datacurators.nl/tide/). PCR primers used for TIDE/ICE genotyping are listed in **Supplementary Table 4.**

### Cell viability assay

Cells were incubated with varying concentrations of DLK1-ADC or isotype ADC and cultured for 72 hours. Cell viability was determined as the percentage of live cells based on forward and side scatter and/or propidium iodide negativity via flow cytometry.

### ClickiT Edu cell proliferation assay

Cell proliferation was quantitated using the Click-IT EdU cell proliferation assay (Invitrogen, Waltham, MA). Cells were incubated with Click-IT EdU dye for 4 hours and samples were acquired on Quanteon flow cytometer.

### ChIP-seq

ChiP-seq was performed as previously described (61). Chromatin from human CD34+ HSPCs and from *in vitro* differentiated megakaryocytes and monocytes (as described above) was immunoprecipitated with H3K4me3 (Abcam, Cambridge, United Kingdom), H2K27ac, and H3K27me3 (both Diagnode, Seraing, Belgium). Library preparation, sequencing and bioinformatic analysis followed the published protocol except that the reads were aligned to human genome 19.

### CUT&RUN

Patient leukemic blasts were sorted by FACS. Blasts were defined as DAPI^-^, CD45^dim^ and CD33^+,^ with lineage exclusion CD3^-^/CD7^-^/CD19^-^. Per antibody, 250,000 cells were sorted. CMK cells expressing doxycycline-inducible HA-GATA1s were harvested 48h after doxycycline induction. CUT&RUN was performed as previously described (50, 62, 63). CMK cells 100,000 cells per condition were sorted and profiled for rabbit anti-H3K4me3 (EpiCypher, Durham, NC), rabbit anti-H3K27ac (Diagnode), rabbit anti-H3K27me3, and HA specific antibody (both Cell signaling technologies).

The pAG/MNase nuclease (Addgene #123461) was produced and purified as previously described (63) after removal of the HA tag. Sequencing was performed on an Illumina NextSeq machine at the sequencing core of the Max Planck Institute for Molecular Genetics (MPIMG, Berlin). Paired, raw reads were demultiplexed, resulting in reads 42 bases in length. The reads were quality- and adapter trimmed using BBduk, then aligned to the human reference genome (hg19), using BWA (Version: 0.7.17-r1188) in ‘aln’ mode.

### CUT&TAG

CUT&TAG was performed as described in the EpiCypher CUTANA protocol v1.5 with minor adaptations. For each sample, 350,000 viable blasts were sorted as described above. After nuclear extraction, 10,000 nuclei were allocated per antibody condition. Primary antibody incubation was carried out in 0,01% Digitonin buffer. The pAG-Tn5 was obtained from Epicypher. Libraries were generated using NEBNext® High-Fidelity 2X PCR Master Mix (New England Biolabs) with barcoded i5/i7 primers (Integrated DNA Technologies, Leuven, Belgium).

### ATAC-seq

For each library, 50,000 leukemic blasts were FACS-sorted as described above. Nuclei were prepared and tagmented according to the manufacturer’s instructions using the Tagment DNA enzyme and Buffer kit (Illumina, San Diego, CA). Transposed DNA was purified with MinElute PCR Purification Kit (Qiagen). Libraries were generated using NEBNext® High-Fidelity 2X PCR Master Mix (New England Biolabs) with barcoded i5/i7 primers (Integrated DNA Technologies).

### Methocult colony forming assay

1000 cells were mixed with Isotype ADC or DLK1-ADC and resuspended in Methocult containing media. The cell suspension was plated in a 6-well plate and incubated in a 37°C CO_2_ incubator. The number of colonies were counted on day 10.

### *In vivo* studies

NSG-SGM3 mice (Stock#010636) purchased from Jackson Laboratories (Bar Harbor, ME, USA) were transplanted with leukemic cells (1.0 - 2.5 × 10^6^) by tail vein injections. For competitive homing assay, Violet proliferation dye stained CMK cells were co-injected with CFSE stained *DLK1* KO cells (10^6^ each) via the tail vein and mice were euthanized 72 hours after. The percentage of CMK and DLK1 KO cells in the bone marrow was determined post euthanasia by flow cytometry as described previously (64). Mice were maintained in the Nemours Life Science Center following the guidelines established by the Nemours IACUC. Following engraftment confirmation, mice were either left untreated or treated with indicated doses of DLK1-ADC or Isotype ADC via the tail vein. The leukemic burden was measured by flow cytometry of mouse peripheral blood drawn by submandibular bleeds as described previously (65, 66). Mice were monitored daily, and once mice reached experimental endpoints they were euthanized. Immunohistochemistry was performed as described previously (64).

### Protein-expression

Body-wide protein expression for DLK1 was summarized from the Human Protein Atlas (HPA) (REF: HPA). Data was retrieved for DLK1 (HGNC: DLK1) from HPA v24.0, release date September 22, 2024 and plotted without re-scaling.

### Ethics statement

All PDX samples, patient samples and primary human CD34⁺ cells were obtained with informed consent under protocols approved by the Institutional Review Boards of the contributing institutions (Nemours Children’s Health, Goethe University Frankfurt and collaborating centers).

### Statistical analysis

Statistical evaluations of experimental data were carried out using 2-way analysis of variance (ANOVA) or unpaired t-test was used for comparisons between two groups. To calculate the survival of mice the Kaplan-Meier survival estimate was used. A value of p < .05 was considered statistically significant. All data are presented as standard error of the mean (SEM). Calculations were performed using GraphPad Prism 10 (STATCON). Statistical analyses of gene expression data (RNA-seq) were carried out in R using edgeR or via msigdr.

## Author Contributions and Disclosures

LV and SPB designed and performed experiments, analyzed data and wrote the manuscript. MT and JRF designed and performed experiments and collected and analyzed data for some of the *in vitro* studies. JG-D, KS, RB, AS, RER, SM collected and analyzed data. JH and YP provided some of the PDX models used in the study. PHvB provided antibody drug conjugates. PHvB, JH, AS, YP, SP, EAK edited the manuscript. J-HK designed the study, supervised the research, analyzed the data and wrote the manuscript. AG designed the research study, performed experiments, analyzed the data, supervised the research and wrote the manuscript. J-HK has advisory roles for Boehringer, Roche and Jazz Pharmaceuticals. Patrick van Berkel is employed by ADC Therapeutics. None of the other authors has a relevant conflict of interest to disclose.

## Acknowledgements

Funding from Lisa Dean Moseley Foundation, Leukemia Research Foundation of Delaware, Department of Defense W81XWH-22-1-0981, and Nemours Foundation is gratefully acknowledged. The study was supported by grants to JHK from the German Research Foundation (DFG; KL 2374/6-1) and *Hilfe für Krebskranke Kinder e.V.* as part of the C^3^OMBAT-AML research units. RB is a fellow of the Mildred Scheel Career Center (MSNZ). DH was supported by the Frankfurt Foundation for Children with Cancer. The use of Cell Science Core, Histochemistry and Tissue Processing Core, and Biobank and Molecular Analysis Program partially supported by the Delaware INBRE (GM103446) is acknowledged. The RNA sequencing of Down syndrome AML was funded by Target Pediatric AML (TpAML: https://targetpediatricaml.org/). Work was also performed under the auspices of the U.S. Department of Energy by Lawrence Livermore National Laboratory under contract DE-AC52-07NA27344. We thank Skyler Brand and Sai Lasya Agasthya Reddy for technical assistance.

