## Supplementary for "DLK1 is a GATA1s-Driven Dependency and Therapeutic Target in Down Syndrome-Associated Myeloid Leukemia"

### Supplementary Figures

A

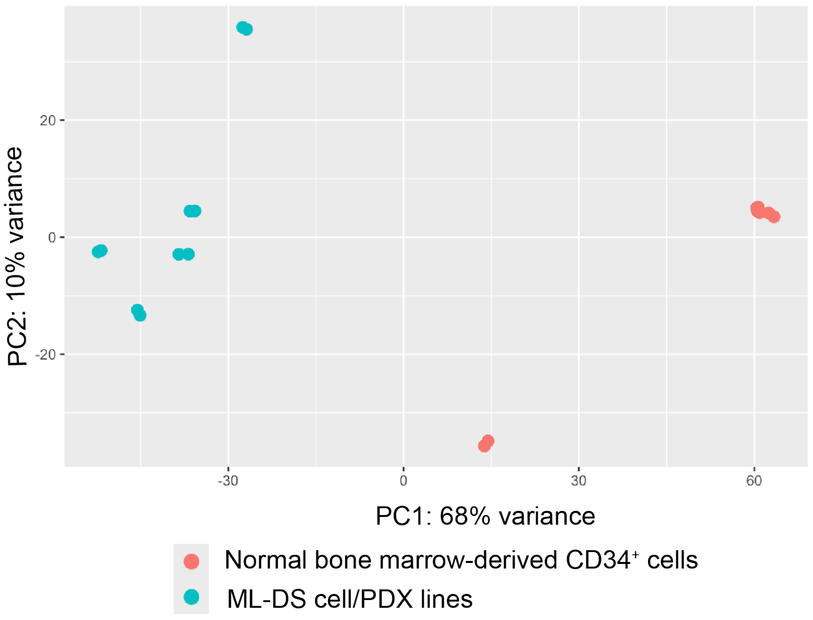


B

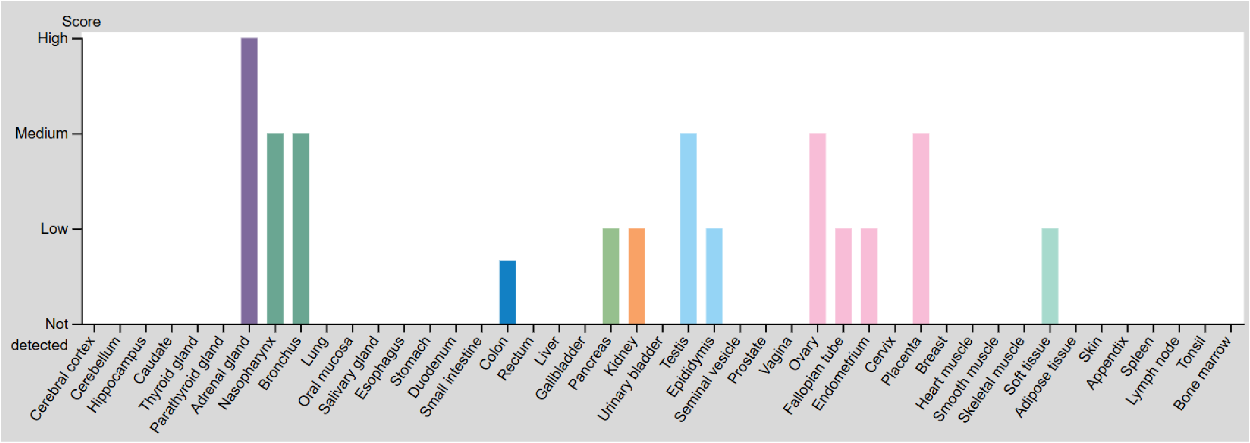


**Supplementary Figure 1**

**A**) Principal component analysis plot showing the clustering of ML-DS cell line/PDX samples (black) and $CD34^+$ samples (red) in RNA-seq analysis.

**B**) DLK1 expression score from the Protein atlas.

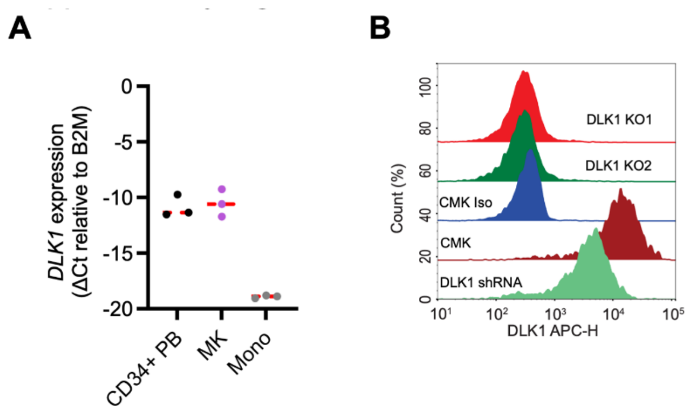


**Supplementary Figure S2.**

**A**) qRT-PCR for DLK1 expression relative to B2M.

**B**) Flow cytometry plot showing the cell surface expression of DLK1 in CMK *DLK1* shRNA and *DLK1* knockout clones.

### Supplementary Tables

**Supplementary Table 1**
Differentially regulated genes encoding cell surface resident proteins in ML-DS samples compared to normal samples. Genes with adjusted pvalue <0.05 were selected and rank ordered based on log2 fold change.

**Supplementary Table 2: qRT-PCR primers and Taqman assays**

| Target gene | Direction | Sequence |
|---|---|---|
| DLK1 | Forward | AGGGTCCCCTTTGTGACCA |
| | Reverse | GCAGGCCCGAACATCTCTATC |
| GAPDH | Forward | GCTGTCCAACCACATCTCCTC |
| | Reverse | TGGGGCCGAAGATCCTGTT |
| DLK1 | Taqman | Assay ID: Hs00171584_m1 |
| MEG3 | Taqman | Assay ID: Hs00292028_m1 |
| B2M primer-limited | Taqman | Assay ID: Hs99999907_m1 |

**Supplementary Table 3: sgRNA and shRNA name, system, and spacer sequences**

| sgRNA name | System | Spacer sequence |
|---|---|---|
| sgLuc2 | CRISPR-Cas9 | GTCCCAAAACAACAACGGCGGC |
| sgRNA A#1 | CRISPR-Cas9 | GGCCCCCGGCAGCTGGGCAG |
| sgRNA A#2 | CRISPR-Cas9 | GAGCAGGGGACGCTGAGCTG |
| sgRNA B | CRISPR-Cas9 | GGCGTGGGGCTGCTGGGGCA |
| sgRNA#1 | CRISPR-Cas9/RNP | GGTCACGCACTGGTCACAAA |
| sgRNA#2 | CRISPR-Cas9 | gAGTCCTTTCCCGAGTACCCG |
| shRNA | shRNA | GGTGTCCATGAAAGAGCTC |

**Supplementary Table 4: PCR primers**

| Target | Direction | Sequence | Purpose |
|---|---|---|---|
| DLK1 exon 3 | Forward | TCTCTACCCCTGCCCTCTTC | TIDE, lenti |
| | Reverse | ATCACTGGGCTTCCATGACC | TIDE, lenti |
| DLK1 exon 3 | Forward | CCTGTCTGCGTTCTCAGAGG | TIDE, RNP |
| | Reverse | GGTTCTCCACAGAGTCCGTG | TIDE, RNP |
| DLK1 exon 4 | Forward | CTCAGCTAGGCACGCTGTTG | TIDE, lenti |
| | Reverse | CTCACACGCATCGGGACTT | TIDE, lenti |
| sgRNA A#1 Enh | Forward | CATGCCTAGCCTGGAGGAAG | TIDE, lenti |
| | Reverse | ACCCAGCGTCACCAACATAG | TIDE, lenti |
| sgRNA B Enh | Forward | TTGTGGTAAAGGCCACGCTGTT | TIDE, lenti |
| | Reverse | ACGTCCCTCTCTCCTGAAAGCAT | TIDE, lenti |
